# Differential DNA methylation in blood in nuclear genes that encode mitochondrial proteins in mild cognitive impairment and Alzheimer’s disease

**DOI:** 10.1101/2025.06.16.659556

**Authors:** Alexander E. Boruch, Andy Madrid, Ligia A. Papale, Phillip E. Bergmann, Isabelle Renteria, Samuel Faasen, Dane B. Cook, Reid S. Alisch, Kirk J. Hogan

## Abstract

**Background:** Altered mitochondrial function contributes to the pathogenesis of mild cognitive impairment (MCI) and late-onset dementia due to Alzheimer’s disease (AD).

**Objective:** To test for differential methylation in nuclear genes that encode proteins that participate in mitochondrial function between cognitively unimpaired participants (CU) and those with MCI and AD.

**Methods:** Recently published whole genome methylation sequencing (WGMS) in blood from CU participants (*N*=174), and those with MCI (*N*=99) and AD (*N*=109) was used to test for differential methylation in 1,121 nuclear genes that encode proteins that participate in mitochondrial function in the *Human Protein Atlas*.

**Results:** Seventy-four nuclear genes that encode proteins that participate in mitochondrial function were differentially methylated between persons with MCI and CU. Seventy-one genes were differentially methylated between persons with AD and CU, and 132 genes were differentially methylated between persons with MCI and AD. Thirteen differentially methylated genes shared between the 3 comparisons support contributions from disrupted metabolism and oxidative stress pathways in AD pathogenesis.

**Conclusions:** Nuclear genes that encode proteins that participate in mitochondrial glucose metabolism, fatty acid metabolism and oxidative stress pathways are differentially methylated between persons who are CU and those with MCI and AD.

## INTRODUCTION

Mitochondrial dysfunction contributes to MCI and AD pathogenesis,^1^ with altered mitochondrial function observed in the brain and blood of persons with dementia.^2–4^ While the full molecular etiology of mitochondrial dysfunction in MCI and AD remains uncertain, evidence indicates that environmental influences increase AD risk through interactions with DNA methylation.^5^ DNA methylation comprises the covalent addition of a methyl group to a nucleotide, primarily to cytosine in CpG dinucleotides, that coordinates gene expression.^6^ Using whole genome methylation sequencing (WGMS) we identified differential DNA methylation in amyloid precursor protein (*APP*), presenilin-1 protein (*PSEN1*) presenilin-2 protein (*PSEN2*), apolipoprotein E (*APOE*), and microtubule associated protein tau (*MAPT*) loci correlated with MCI and AD susceptibility and pathophysiology.^7,8^ We observed differentially methylated genes in persons with MCI and AD in pathways corresponding to accelerated cognitive decline (*e.g*., synaptic membrane integrity, regulation of neuron projection development, and cation channel complexes),^9^ thereby demonstrating that DNA methylation levels in accessible peripheral tissues carry the potential for improved MCI and AD diagnosis and prognosis.^10,11^ Multiple nuclear genes that encode proteins that participate in mitochondrial function are downregulated in blood from persons with MCI.^12^ Accordingly, we used whole genome methylation sequencing (WGMS) in CU participants and in participants with MCI and AD to identify differentially methylated nuclear genes that encode mitochondrial proteins in *The Human Protein Atlas* (N=1,121 genes).^13^

## MATERIALS AND METHODS

### Study Populations

This research was conducted in accord with the Declaration of Helsinki. The experimental protocol was approved by the institutional review board (IRB) of the University of Wisconsin School of Medicine and Public Health, Madison, WI (Health Sciences IRB #2015-0030 & #2023-1522). Written consent for study participation was obtained from each participant. Participants were enrolled in the Wisconsin Alzheimer’s Disease Research Center (WADRC),^14^ and the Wisconsin Registry for Alzheimer’s Prevention (WRAP).^15^ Details of the study design and demographic characteristics have been previously published.^8,9^ Participants were classified as cognitively unimpaired (CU) or as meeting criteria for MCI or AD based on based on the National Institute on Aging-Alzheimer’s Association thresholds by consensus conference.^16,17^ Samples from participants whose clinical status reverted to CU, or who were diagnosed as having non-Alzheimer’s disease dementia at a subsequent visit, were excluded.

### Whole genome methylation sequencing (WGMS): DNA Extraction, Data Processing, and Statistical Analysis

Detailed methods of blood samples acquisition, whole genome methylation sequencing (WGMS), data processing, and statistical analyses have been described in our prior work. In brief, high molecular weight genomic DNA was obtained from blood samples acquired on the visit nearest the date of MCI and AD diagnosis and age-matched to CU cohort samples. Sequence libraries were constructed using the NEBNext Enzymatic Methyl-seq (EM-seq™) kit (Ipswitch, MA) for whole genome methylation sequencing on an Illumina NovaSeq6000 sequencer (San Diego, CA). Image processing and sequence extraction used the Illumina Pipeline (Illumina, San Diego, CA), and raw WGMS data were aligned to the GRCh38.14 (hg38) human reference sequence.

For 2-way cognitive status comparisons (*i.e*., MCI *vs*. CU, AD *vs*. MCI, and AD *vs*. CU) a CpG locus with a LFDR < 0.05 and an estimated methylation difference ≥ 2.5% was used to identify a differentially methylated position (DMP). DMPs were annotated to genomic structures with *annotatr* (version 1.28.0)^18^ for all known isoforms. Differentially methylated genes (*i.e.,* DNA sequences spanning 3 kilobases (kb) 5′ of a transcription start site (TSS) to 200 base pairs (bp) 3′ of a transcription termination site (TTS)) were defined as those that comprise a gene-wide LFDR < 0.01 between groups that differ by cognitive status and contain at least one DMP. Gene promoters (*i.e.,* regions within 5 kilobases (kb) in the 5’ direction and 200 base pairs (bp) in the 3’ direction of a transcription start site (TSS)) and known enhancers of nuclear genes encoding proteins with mitochondrial function that contained at least one DMP were also extracted and characterized.

### Identification of differentially methylated genes that encode proteins that participate in mitochondrial function

The *Human Protein Atlas* comprises a comprehensive inventory of the human proteome comprising all cell and tissue types assembled with an integrated-omics approach including antibody-based imaging, mass spectrometry-based proteomics and transcriptomics. The *Human Protein Atlas* has supported multiple investigations of genome-disease interactions including in AD.^13,19–21^ Nuclear genes that encode proteins with mitochondrial functions selected for investigation were exclusively located within the outer mitochondrial membrane, the inner mitochondrial membrane and the matrix of mitochondria, or were located within mitochondria in addition to other cellular organelles and structures. R package *dplyr* (version 1.1.4) was used to test for intersections between the *Human Protein Atlas* database and differentially methylated nuclear genes identified in each disease pairwise comparison (*i.e.,* MCI *vs.* CU, AD *vs.* CU, AD *vs.* MCI).^9,22^

### Statistical analysis and visualization software

Analyses were performed in *R-Studio* (version 23.12.1+402). Histograms were created with *ggplot2* (version 3.5.2).^23^ Venn diagrams of differentially methylated genes shared between comparisons were generated using R package *eulerr* (version 7.0.2).^24^ Bar plots and pie charts were generated using *GraphPad Prism* (version 10.2.2.).

## RESULTS

### Differential DNA methylation in nuclear genes encoding proteins with mitochondrial functions between persons with MCI and CU

One hundred and eighty DMPs were identified in nuclear genes that encode proteins with mitochondrial functions between MCI and CU participants (Supplemental Table S1A). The DMPs were distributed in all gene structures with 88.89% in introns and 26.67% in promoters (Figure 1). A majority of DMPs (125/180) were hypermethylated (*i.e.*, greater DNA methylation in MCI compared to CU participants) with a maximum methylation difference of 6.3% (Figures 2A & 2B). The remaining DMPs (54/180) were hypomethylated (*i.e.*, lower DNA methylation in MCI compared to CU participants) with a maximum methylation difference of −6.5%. Analyses of promoter-enhancer interactions identified 3 promoter regions comprising one or more DMPs that interact with 12 distinct enhancer regions and 3 enhancer regions with one or more DMPs that interact with 2 unique gene promoters in participants with MCI (Supplemental Tables S2A & S2B).

**Figure 1:**
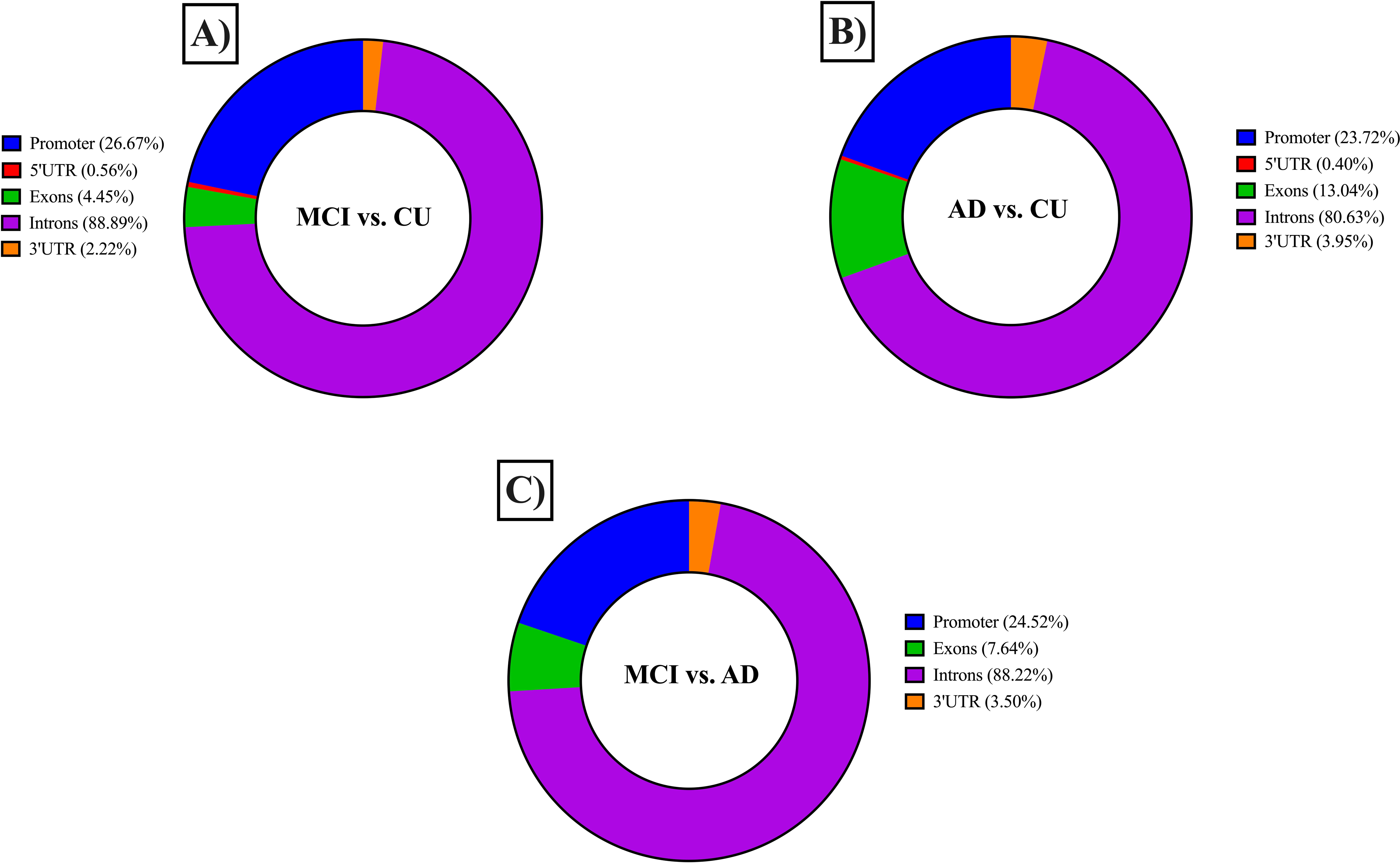
Distributions of differentially methylated positions (DMPs) relative to genomic structures within nuclear genes that encode mitochondrial pathway proteins. DMPs were distributed in gene structures in MCI *vs*. CU (A), AD *vs*. CU (B), and AD *vs*. MCI (C) pairwise comparisons of cognitive status. Promoter regions displayed above encompass the annotatr-defined promoter region (*i.e.,* <1kB DNA sequence upstream of the transcription start site) and the 1-5 kB region upstream of the promoter. Because we defined promoter regions as up to 5 kB in the 5’ direction of the transcription start site, we combined the annotatr-defined promoter region and 1-5 kB regions upstream of the promoter for the purpose of consistency.

**Figure 2:**
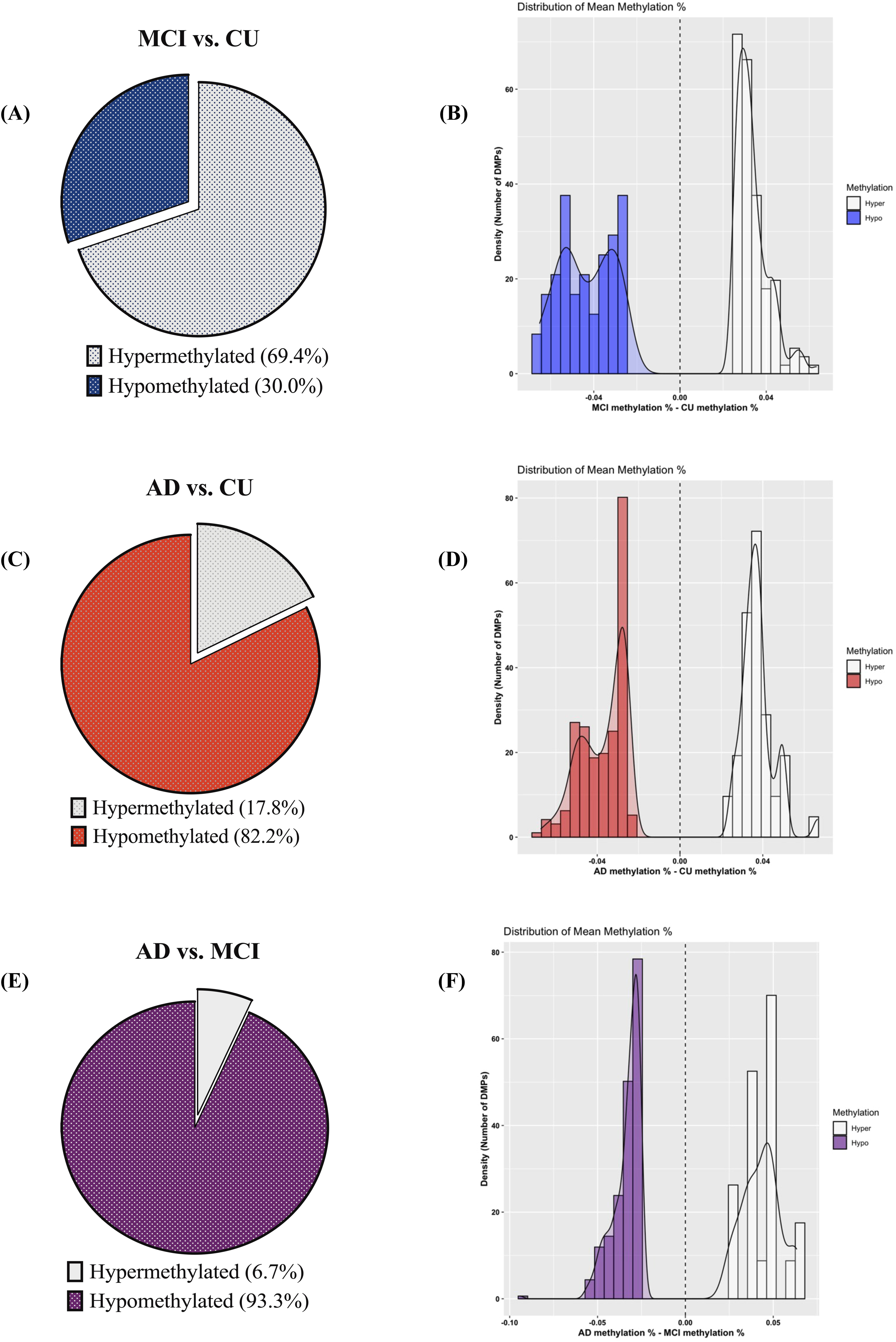
Polarity proportions and mean density of differentially methylated positions (DMPs). Pie charts display the proportion of hypermethylated and hypomethylated DMPs in nuclear genes that encode mitochondrial pathway proteins in pairwise comparisons of **A)** MCI *vs*. CU (180 DMPs; hypermethylated [gray]; hypomethylated [blue]), **C)** AD *vs*. CU (253 DMPs; hypermethylated [gray]; hypomethylated [red]), and **E)** AD *vs*. MCI (314 DMPs; hypermethylated [gray]; hypomethylated [purple]). Density plots depict distributions of the mean methylation differences across DMPs in nuclear genes that encode mitochondrial pathway proteins in pairwise comparisons of **B)** MCI *vs*. CU comparison (blue bars depict hypomethylated DMPs; white bars depict hypermethylated DMPs), **D)** AD *vs*. CU comparison (red bars depict hypomethylated DMPs; white bars depict hypermethylated DMPs), and **F)** AD *vs*. MCI comparison (purple bars depict hypomethylated DMPs; white bars depict hypermethylated DMPs).

Seventy-four differentially methylated nuclear genes that encode proteins with mitochondrial functions were identified between participants with MCI and CU participants (Supplemental Table S3). Differentially methylated genes include hexokinase 1 (*HK1*), hexokinase domain containing 1 (*HKDC1*), glutamate dehydrogenase 1 (*GLUD1*), and hydroxyacyl-CoA dehydrogenase (*HADH*).

### Differential DNA methylation in nuclear genes encoding proteins with mitochondrial functions between persons with AD and CU

Two hundred fifty-three DMPs were identified in nuclear genes that encode proteins with mitochondrial functions between AD and CU participants (Supplemental Table S1B). The DMPs were distributed in all gene structures with 80.63% in introns and 23.72% in promoters (Figure 1). A minority of DMPs (45/253) were hypermethylated (*i.e.*, greater DNA methylation in AD compared to CU participants) with a maximum methylation difference of 6.7% (Figures 2C and 2D). The remaining DMPs (208/253) were hypomethylated (*i.e.*, lower DNA methylation in AD compared to CU participants) with a maximum methylation difference of −6.7%. Analyses of promoter-enhancer interactions identified 3 promoter regions comprising one or more DMPs that interact with 23 distinct enhancer regions and 9 enhancers regions comprising one or more DMPs that interact with 9 unique promoter regions in participants with AD (Supplemental Tables S2A & S2B).

Annotation of these DMPs to genes identified 71 differentially methylated nuclear genes that encode proteins with mitochondrial functions, including acyl-CoA thioesterase 1 (*ACOT1*), acyl-CoA thioesterase 8 (*ACOT8*) and 3-hydroxyisobutyryl-CoA hydrolase (*HIBCH*; Supplemental Table S3).

### Differential DNA methylation in nuclear genes encoding proteins with mitochondrial functions between persons with AD and MCI

Three hundred and fourteen DMPs were identified in nuclear genes that encode proteins with mitochondrial functions between AD and MCI participants (Supplemental File S1C). The DMPs were distributed in all gene structures except the 5’UTR, with 88.22% in introns and 24.52% in promoters (Figure 1). Few DMPs (21/314) were hypermethylated (*i.e.*, greater DNA methylation in AD compared to MCI participants) with a maximum methylation difference of 6.3%. The majority of DMPs (293/314) were hypomethylated (*i.e.*, lower DNA methylation in AD compared to MCI participants) with maximum methylation difference of −9.4% (Figures 2E & 2F). Analyses of promoter-enhancer interactions identified 11 promoter regions with one or more DMPs that interact with 144 distinct enhancer regions and 15 enhancers regions with one or more DMPs that interact with 12 unique promoter regions in participants with AD compared to MCI (Supplemental Tables S2A & S2B).

Annotation of these DMPs to genes identified 133 differentially methylated nuclear genes that encode proteins with mitochondrial functions that participate in responses to oxidative stress, including BCL2 apoptosis regulator (*BCL2*), TNF receptor associated factor 6 (*TRAF6*), heat shock protein family A (Hsp70) member 9 (*HSPA9*), heat shock protein family D (Hsp60) member 1 (*HSPD1*), and mitochondrial calcium uniporter dominant negative subunit beta (*MCUB*; Supplemental Table S3).

### Differential DNA methylation in nuclear genes encoding proteins with mitochondrial functions shared between MCI vs. CU, AD vs. CU, and AD vs. MCI pairwise comparisons

Thirteen differentially methylated nuclear genes that encode proteins with mitochondrial functions were shared between the 3 pairwise comparisons (Figure 3). Three shared genes that encode proteins with mitochondrial functions participate in signal transduction, such as amyloid beta precursor protein binding family B member 2 (*APBB2*), ganglioside induced differentiation associated protein 1 (*GDAP1*) and regulator of G protein signaling 7 (*RGS7*). Hexokinase domain containing 1 (*HKDC1*), which participates in glucose-6-phosphate metabolism, was also shared among the 3 pairwise comparisons. University of California-Santa Cruz Genome Browser^25^ schematic diagrams display DMPs within *APBB2* (Figure 4A), *HKDC1* (Figure 4B), and *RGS7* (Figure 4C) for each of the 3 pairwise comparisons.

**Figure 3:**
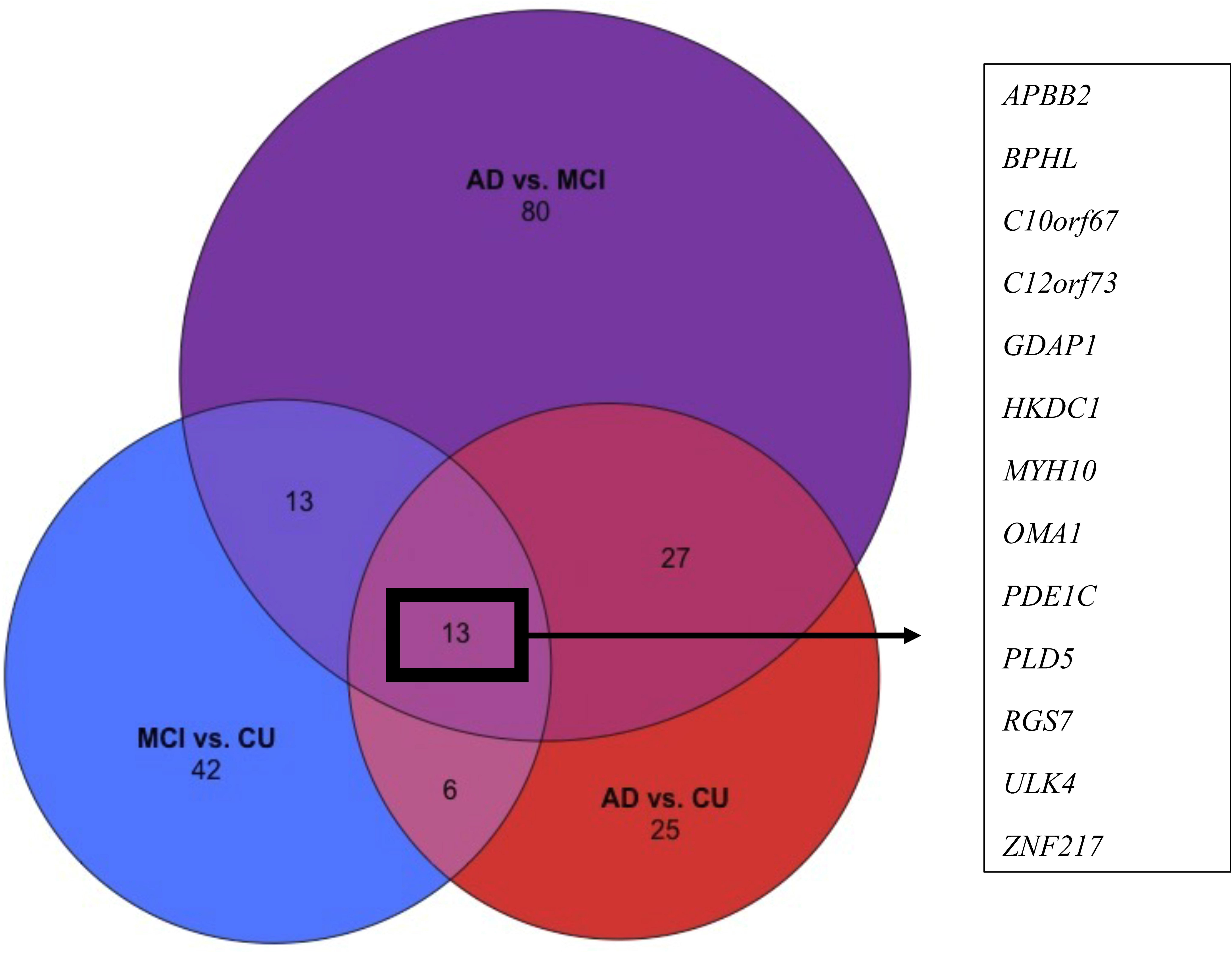
Differentially methylated genes shared between pairwise comparisons. A 3-way Venn diagram showing the shared DMPs between MCI *vs*. CU (blue, *N*=74), AD *vs*. CU (red, *N*=71), and AD *vs*. MCI (purple, *N*=133) pairwise comparisons of cognitive status. The gene symbols for the 13 differentially methylated genes shared between the 3 pairwise comparisons are shown.

**Figure 4:**
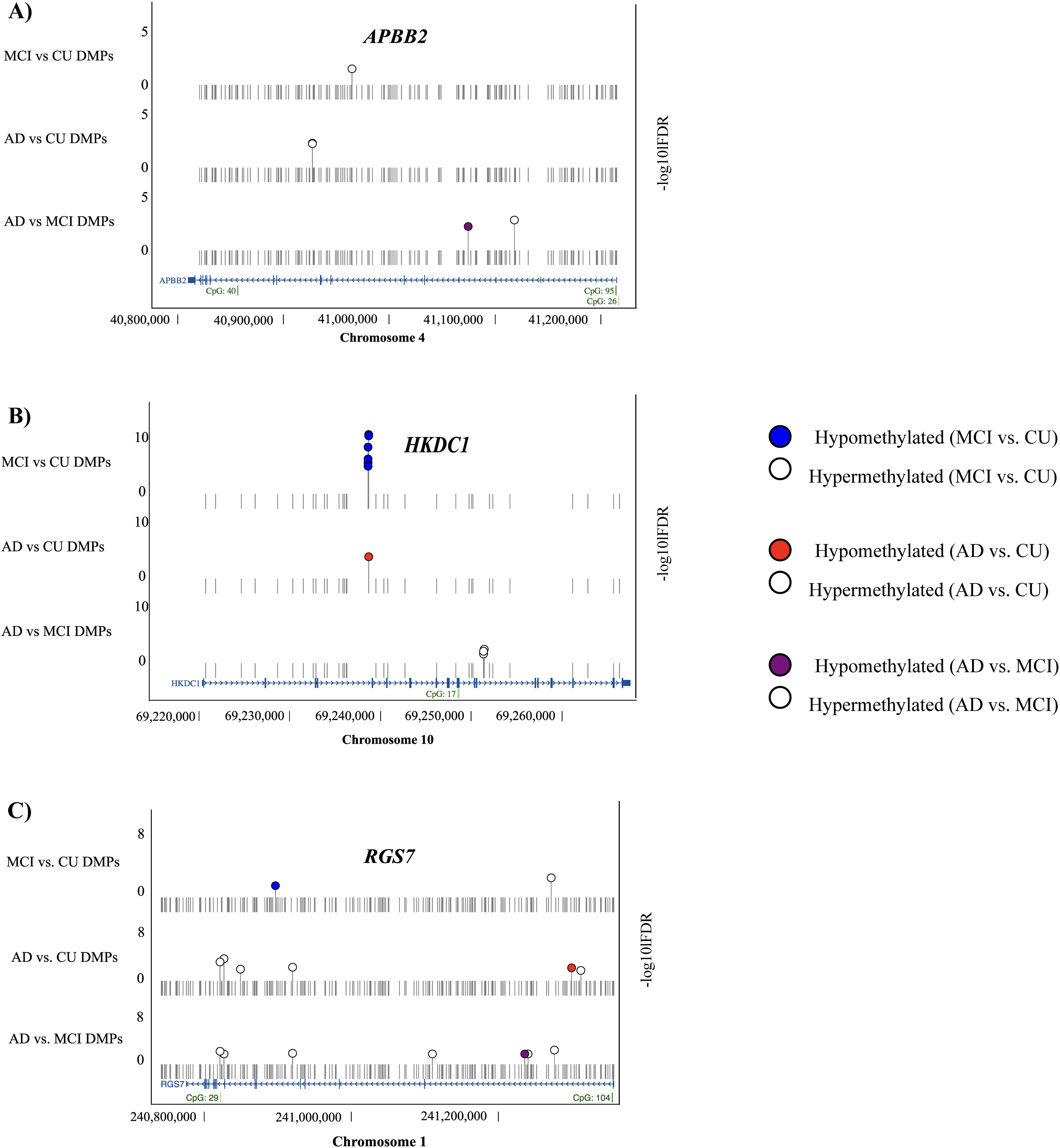
Schematic diagrams of differentially methylated nuclear genes (DMGs) that encode mitochondrial pathway proteins shared between persons with and without mild cognitive impairment (MCI) and Alzheimer’s disease (AD). (A)–(C). Sense (>) and anti-sense (<) strands are depicted with gene name abbreviations above each panel. Alignment to human reference genome hg38 coordinates are displayed in base pairs (bp) at the bottom of each panel with the chromosome number (*x*-axis). The direction of gene transcription is depicted by small arrowheads on the blue line. Blue rectangles indicate coding exons connected by a blue line that indicate introns. CpG islands > 300 bp in length are indicated by green rectangles. The relative location of CpG dinucleotides in each gene is shown as a vertical line with circles on top, and one in 25 non-significant CpGs in each gene are shown (gray vertical line with circle, N.S.). The significance level (*y* axis) of differentially methylated CpGs is shown for MCI versus CU (top row; blue circles are hypomethylated, white circles hypermethylated), AD versus CU (middle row; red circles are hypomethylated, white circles hypermethylated), and AD versus MCI (bottom row; purple circles are hypomethylated, white circles hypermethylated). A corrected significance level of local false discovery rate < 0.05 and > 2.5% differential methylation level was adopted for all comparisons.

## DISCUSSION

Mitochondrial dysfunction contributes to MCI and AD pathogenesis.^1,26,27^ Present data provides evidence of widespread differential DNA methylation of nuclear genes that participate in diverse mitochondrial pathways in MCI and AD. The differentially methylated genes from the MCI *vs*. CU pairwise comparison displayed DMPs with overall hypermethylation (69.4%) involved in mitochondrial pathways, while the AD *vs*. CU and AD *vs*. MCI pairwise comparisons both exhibited DMPs with overall hypomethylation. These differential DNA methylation trends in nuclear encoded with mitochondrial functions between persons who differ by cognitive status are consistent with our previously published WGMS findings^9^ and in accord with reductions in global methylation levels that occur with normative aging.^28^

Hypometabolism and impaired glucose uptake are observed in the peripheral and central nervous systems of persons with MCI.^29–31^ Glucose dysregulation develops years in advance of cognitive decline in correlation with changes in cognitive status and blood brain barrier disruption.^32,33^ Differential methylation of *HKDC1*, a protein-coding gene in the hexokinase family that catalyzes glucose conversion to pyruvate, participates in the pathogenesis of hypometabolism (*e.g.,* hyper-insulinemic hypoglycemia).^34–36^ Hyper-insulinemic responses are associated with mutations in *HK1*, *GLUD1*, and *HADH*, all of which are differentially methylated between participants with MCI and those who are CU.^37,38^ In turn, chronic hyperinsulinemia increases the risk neurodegenerative diseases including AD.^39,40^

Neuronal tissue depends upon glucose as a primary fuel source and ketone bodies as a secondary fuel source that accounts for high oxygen demand with the potential for generating reactive oxygen species that arise from beta oxidation.^41^ Fatty acid permeation through the blood-brain barrier,^42^ and differences in plasma fatty acid profiles of persons with AD,^43^ link markers of beta oxidation to AD clinical outcomes.^44^ Reactive oxygen species disrupt intracellular calcium regulation and promote amyloid plaque accumulation in persons with AD.^45^ Multiple mitochondrial proteins encoded by nuclear genes that are differentially methylated between AD participants and those with CU facilitate beta-oxidation (*e.g.,* acyl-CoA thioesterase 1 [*ACOT1*] and 8 [*ACOT8*]), amyloid beta accumulation (*e.g.,* amyloid beta precursor protein binding family B member 2 [*APBB2*]), and calcium regulation (*e.g.,* adenylate cyclase [*ADCY1*]). Differential DNA methylation of nuclear genes that encode mitochondrial beta-oxidation proteins may alter responses to suppressed long-chain fatty acid hydrolysis (*ACOT1*) and oxidation (*ACOT8*). Differential DNA methylation of nuclear genes related to amyloid beta (*APBB2*) and calcium regulation (*ADCY1*) aligns with AD mechanisms reported in murine models with widespread neuronal death.^46^

Oxidative stress generated by reactive oxygen species and free-radical byproducts contributes to the pathogenesis of both MCI and AD^47,48^ eliciting DNA damage, impaired calcium regulation, and neuronal apoptosis.^49,50^ DMPs within genes encoding oxidative stress-related proteins were identified in pairwise comparisons between persons with AD and MCI. Proteins encoded by nuclear genes that are differentially methylated between participants with AD and MCI also contribute to cellular distress responses (*e.g.,* heat shock protein family A member 9 [*HSPA9*] and heat shock protein family D member 1 [*HSPD1*]), and innate immune cell mobilization (*e.g.,* TNF receptor associated factor 6 [*TRAF6*]). Proteins encoded by nuclear genes that are differentially methylated between participants with AD and MCI play central roles calcium regulation (*e.g.,* [*MCUB*]) and apoptotic immune cell death (*e.g.,* BCL2 apoptosis regulator [*BCL2*] and BCL2 like 13 [*BCL2L13*]) and may engender systemic distress responses. Interactions between peripheral and central inflammation in both MCI and AD is mobilized in part through oxidative damage.^50,51^

Thirteen differentially methylated nuclear genes that encode proteins that participate in mitochondrial functions are shared between the 3 pairwise comparisons (Figure 3). Three differentially methylated nuclear genes encode signal transduction proteins. Glucose hypometabolism secondary to toxic levels of glutamate and calcium signal transducers in neurons contributes to declining cognitive status.^52^ The GDAP1 protein participates calcium homeostasis, and *GDAP1* knock-out animal models exhibit interactions between both calcium dysregulation and hypometabolism via pyruvate dehydrogenase inhibition.^53^ Amyloid plaques associated with APBB2, a protein that interacts with the cytoplasmic domains of amyloid beta precursor protein (APP) and amyloid beta precursor-like protein 2, exacerbate synaptic glutamate levels and contribute to neurotoxicity.^52^ Differential DNA methylation within genes that encode proteins participating in mitochondrial functions suggests that chronically reduced metabolic function (*e.g.*, hypometabolism) and associated oxidative stress may partially contribute to AD pathogenesis through reduced neuronal integrity.

### Strengths & Limitations

Strengths of the present study include a large sample size, well-characterized cognitive status phenotypes based on both clinical and psychometric criteria, use of comprehensive WGMS to identify CpG candidates for differential methylation unconstrained by limited microarray CpG coverage, and nucleotide level DMP resolution using enzyme-based rather than bisulfite-based cytosine conversion. A further strength is use of an easily accessible peripheral tissue in keeping with shared DNA methylation alterations observed in blood and brain samples from persons with AD.^54^ Limitations include a cohort of limited diversity, use of a cross-sectional rather than longitudinal experimental design, and testing white blood cells in bulk corrected for cell type proportion rather than testing single white blood cell types. Confirming whether nuclear genes that encode glucose metabolism pathway proteins are differentially methylated between MCI and CU, those that encode fatty acid metabolism pathway proteins are differentially methylated between AD and CU, and those that encode oxidative stress pathway proteins are differentially methylated between AD and MCI requires longitudinal samples from participants as the signs and symptoms of AD progress. Further, differential methylation was measured in nuclear DNA rather than mitochondrial DNA in the present investigation. Mitochondrial DNA sequence variations are associated with increased risk for AD,^55^ and reduced cerebral spinal fluid levels of mitochondrial DNA may correspond with neuronal damage.^56^ However, the contribution of differential mitochondrial DNA methylation to the pathogenesis of neurodegeneration is controversial, and if present, unlikely to play a role in the control of mitochondrial function.^57^

## CONCLUSIONS

Nuclear genes that encode proteins that participate in mitochondrial glucose metabolism, fatty acid metabolism, and oxidative stress pathways are differentially methylated between persons who are cognitively unimpaired (CU) and those with mild cognitive impairment (MCI) and Alzheimer’s disease (AD). Marked hypermethylation of nuclear mitochondrial pathway genes characterizes progression from CU to MCI, whereas marked hypomethylation characterizes progression from MCI to AD at these loci. Differentially methylated genes and differentially methylated positions in nuclear genes of mitochondrial pathways may serve as candidate blood biomarkers for MCI and AD onset and progression.

## ACKNOWLEDGMENTS

The authors thank the Wisconsin Registry for Alzheimer’s Prevention (WRAP) and Wisconsin Alzheimer’s Disease Research Center (WADRC) participants and study personnel who made this work possible.

## FUNDING

Funding was provided in part by the National Institutes of Health, Grant/Award Numbers: R01AG066179; R01AG021155; R01AG027161; P30AG062715; HG003747; and S10OD025245; Alzheimer’s Association, Grant/Award Number: AARF19-614533; UW Department of Neurological Surgery; NLM Grant/Award Number 5T15LM007359. AEB was supported by a UW Department of Kinesiology William H. and Virginia F. Harrison Marsh Center Fellowship. The funding sources had no role in the design, collection, analysis and interpretation of the data, writing the report, or decision to submit the article for publication.

## CONFLICT OF INTEREST

The authors have no relevant disclosures.

## DATA AVAILABILITY

The data supporting the findings of this study are openly available within the article and its supplemental material (S1-S4). Full datasets from the overhead project and previously published manuscript are also available within its supplemental material (https://onlinelibrary.wiley.com/doi/abs/10.1002/alz.14474).

## SUPPLEMENTARY MATERIAL

Supplemental Table S1: DMPs

Supplemental Table S2: Promoter-Enhancer Interactions

Supplemental Table S3: Differentially Methylated Genes

## Notes

https://alz-journals.onlinelibrary.wiley.com/doi/10.1002/alz.14474

### Competing Interest Statement

The authors have declared no competing interest.

